# Experimentally Induced Metamorphosis in Axolotl (*Ambystoma mexicanum*) Under Constant Diet Restructures Microbiota Accompanied by Reduced Limb Regenerative Capacity

**DOI:** 10.1101/277285

**Authors:** Turan Demircan, Guvanch Ovezmyradov, Berna Yıldırım, İlknur Keskin, Ayse Elif İlhan, Ece Cana Fesçioğlu, Gürkan Öztürk, Süleyman Yıldırım

## Abstract

Axolotl (*Ambystoma mexicanum*) is a critically endangered salamander species and a model organism for regenerative and developmental biology. Despite life-long neoteny in nature and in captive-bred colonies, metamorphosis of these animals can be experimentally induced by administering Thyroid hormones (THs). However, biological consequences of this experimental procedure, such as host microbiota response and implications for regenerative capacity, remain largely unknown. Here, we systematically compared host bacterial microbiota associated with skin, stomach, gut tissues and fecal samples based on 16S rRNA gene sequences, along with limb regenerative capacity, between neotenic and metamorphic Axolotls. Our results show that distinct bacterial communities inhabit individual organs of Axolotl and undergo substantial restructuring through metamorphosis. Drastic restructuring was observed for skin microbiota, highlighted by a major transition from *Firmicutes*-enriched to *Proteobacteria*-enriched relative abundance and precipitously decreased diversity. Remarkably, shifts in microbiota was accompanied by a steep reduction in limb regenerative capacity. Fecal microbiota of neotenic and metamorphic Axolotl shared relatively higher similarity, suggesting that diet continues to shape microbiota despite fundamental transformations in the host digestive organs. The results provide novel insights into microbiological and regenerative aspects of Axolotl metamorphosis and will establish a baseline for future in-depth studies.

## Introduction

Metazoan genomes have diversified and evolved in the presence of associated host microbiota. The evolution of morphology and function of animal organ systems may have been influenced by interactions with their microbial partners ^1^. From the host perspective, symbioses between metazoans and microbes provide a synergetic impact to operate essential functions for normal growth, development and behavior ^2-4^. Studies on host-microbiome interactions in health and disease conditions indicate that perturbation of the crosstalk between the host and microorganisms may lead to deleterious consequences such as developmental defects ^3,5^, increased susceptibility to infectious diseases ^6,7^ and ultimately fitness costs ^8^. Even though host genotypes ^9^ and environmental factors, such as diet and habitat ^10,11^, were shown to strongly impact the composition and structure of animal microbiota, ecological forces shaping assembly of the host associated microbial communities have still been poorly understood.

Amphibians, which undergo dramatic morphological changes through metamorphosis, exhibit explicitly altered biphasic life stages to tackle developmental challenges. Remarkably, metamorphosis in several marine animal species such as sponges, corals, crabs, sea urchins, an ascidians is mediated by bacterial community ^12^. Thyroid hormones (THs) are key players in initiation and completion of metamorphosis ^13,14^. Natural accumulation (as in anurans) or administration (as in Axolotl) of THs leads to critical reorganization of organs in order to adapt terrestrial life conditions. This adaptive process includes reconstruction or loss of some existing organs and extremities, and formation of new ones ^15,16^. A prominent example of reconstruction is observed in digestive tract of tadpole. In adult frogs acidic stomach and shorter intestine originate from non-acidic stomach and long intestine of tadpole digestive tract ^17,18^. Growing evidence supports the notion that reshaping of organs and composition of microorganisms reciprocally influence each other as functions of bacterial communities are increasingly being linked to host metabolic activities ^19-21^. Hence, unraveling microbiome compositions in various life stages may offer new insights into the life stage-specific microbial patterns.

Axolotl (*Ambystoma mexicanum*), a salamander species of amphibians, possess experimentally validated features, such as high regenerative capacity ^22^, low cancer incidence ^23^, scarless wound healing ^24^, life-long neoteny with the ability to undergo induced metamorphosis ^25^. These characteristics contributed to the recent establishment of Axolotl as a promising vertebrate model organism for regenerative and developmental biology (reviewed in ^26^). Reference resources for this model such as transcriptome ^27,28^ and a draft genome assembly ^29^ are publicly available. Current studies on regeneration in Axolotl have focused on identification of genes, gene networks and pathways activated during limb and tail regeneration by utilizing transcriptome and proteome profiling tools ^16,27,30,31^. Nevertheless, there is very limited data on microbial diversity of salamanders ^32^ and systematic investigation of microbiota diversity in multiple organs of salamander species, particularly that of Axolotl, has been missing in the literature.

In this study, we hypothesized that induction of metamorphosis in Axolotl leads to restructuring of bacterial microbiota due to reorganization of tissues via metamorphosis. We then asked whether such microbiota restructuring is accompanied by changes in the host regenerative capacity. To test the hypothesis and answer the question, we explored the variation in neotenic and metamorphic Axolotl’s microbiota associated with skin, stomach, gut tissues (crypts), and feces using 16S rRNA gene sequencing and compared limb regenerative capacity between two developmental stages. The results provide novel insights into relationship between perturbed microbial communities due to host factors and putative implication of microbiota in regeneration capacity.

## Results

Details of the experimental design were described in the methods section (Fig. 1). Within 2-3 weeks of hormonal treatment of the animals, we observed weight loss, progressive disappearance of the fin and decrease in the gills size; and in approximately two months all animals showed characteristics of accomplished metamorphosis (Fig. 2a). We first performed comparative analysis of microbiota between neotenic and metamorphic Axolotl organs. We then attempted to answer the question whether shifts in microbiota might be accompanied by a change in host regenerative capacity since previous studies provided evidence that limb regenerative capacity is reduced in metamorphic Axolotl ^33^. We reproduced these observations in this study that regeneration is indeed impeded in the limbs of metamorphic Axolotl (Fig. 2b). We observed that all neotenic animals (n=15) regenerated (day 64) the limb in a miniaturized form with four digits. In contrast, slower blastema formation and regeneration process were apparent in metamorphic animals. We continued to observe limb generation in metamorphic Axolotls up to day 150. At day 150, 2 of 15 metamorphic Axolotls (13 %) restored the limb with four digits. Also, 4 of 15 metamorphic animals (27 %) were capable of regenerating amputated limb with 3 digits only. In addition, another 4 animals (27 %) restored limb with only two digits were observed. Rest of the animals (5 out of 15) failed to regenerate a limb to any extent, indicating that the limb generation capacity in metamorphic Axolotls is severely impeded.

**Figure 1.**
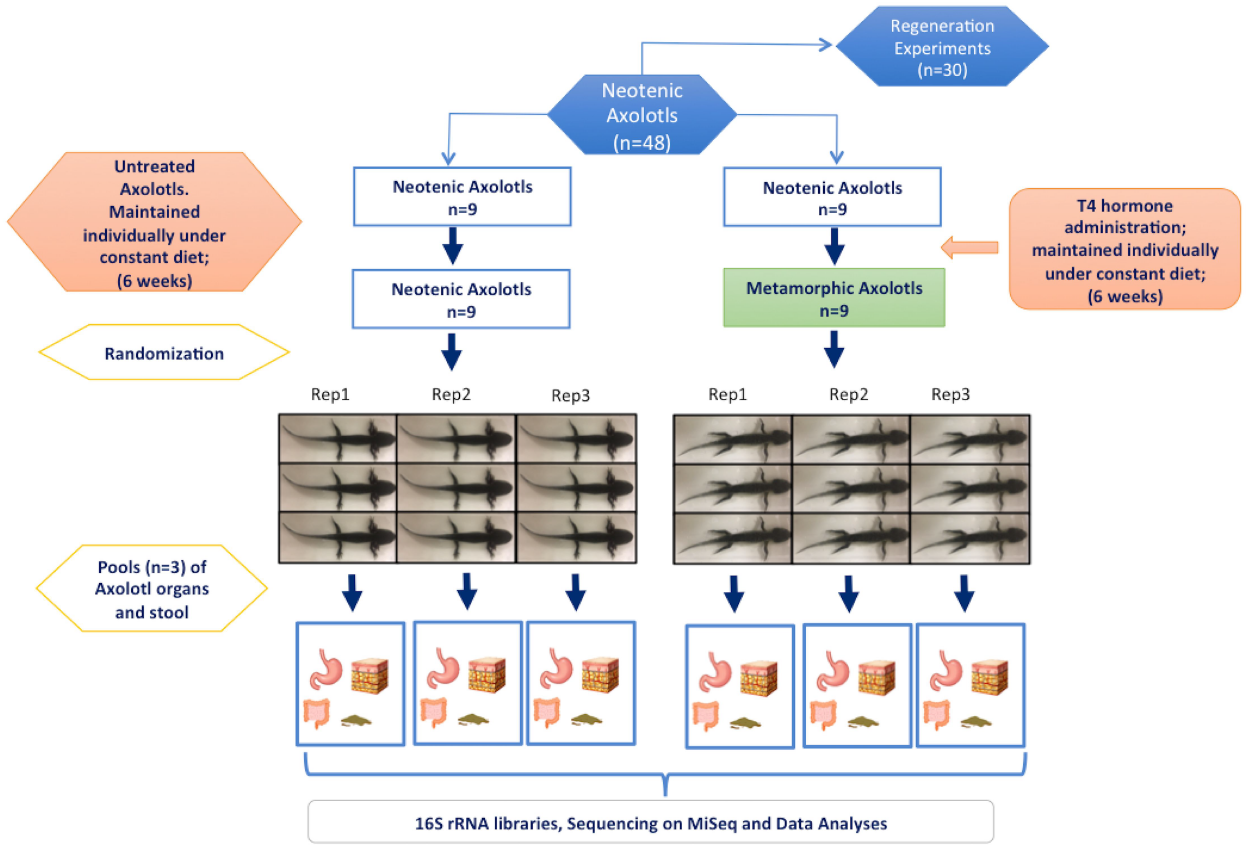
Experimental design for the comparison of neotenic and metamorphic Axolotl. Of 48 siblings, a subset of 24 animals (9 animals for metamorphosis and 15 animals for regeneration experiments) were randomly selected and induced metamorphosis by T4 hormone administration while the rest kept untreated in neoteny. 30 animals (15 neotenic and 15 metamorphosed) were used in regeneration experiments and 18 animals (9 neotenic and 9 metamorphic) were housed for microbiome analysis. Each sample groups for skin, gut, stomach and fecal samples consisted of 3 replicates following randomization and pooling.

**Figure 2.**
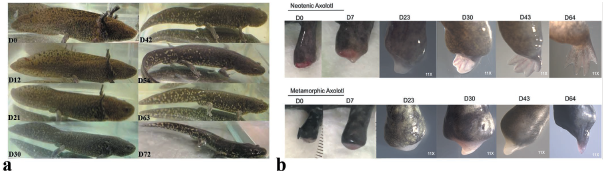
Time course of metamorphosis and limb regeneration. Time course (Day0 - Day72) after T4 administration showing anatomical changes to adapt terrestrial life conditions. Metamorphosis-associated characteristics such as weight loss and disappearance of fin and gills were noticed gradually within this time period. **(a).** Time course (Day0 - Day64) after limb amputation demonstrating differences between regenerative capacity of neotenic (upper panel) and metamorphic (lower panel) Axolotl. Reduction in limb regenerative capacity was observed for metamorphic animals. (n=15 for each group) **(b)**

### Bacterial community structure and membership differ between neotenic and metamorphic Axolotl

Sequencing of the V3-V4 region of the 16S rRNA gene produced approximately 3.7M reads generated from 27 samples (24 samples from Axolotls and 3 aquarium water samples, hereinafter “Aqua”). The sequences were clustered into 14451 high quality, singleton, chloroplast, and mitochondria removed and chimera-checked Operational Taxonomic Units (OTUs). Average amplicon sequences per sample was 139059 ± 49159 sequences). Our data included 12224 (85% of total) *de novo* OTUs (OTU IDs that begin with “New.Reference” or “New.Cleanup.Reference”). We then classified a representative sequence of these OTUs using RDP classifier (v. 2.2) at 70% bootstrap cutoff. We identified 621 OTUs that did not find hits in the RDP database even at the phylum level (“Unclassified Bacteria”). We thus used MOLE-BLAST to determine their identity. Except a few high abundance OTUs (~5% abundance) enriched in the stomach samples hitting plant mitochondria (discarded), most of these OTUs had abundance below 0.1%, which can be considered a rare OTU ^34^. The majority of these OTUs were phylogenetically related to the phylum *Proteobacteria* or *Verrucobacteria* (Supplementary Fig. S1; NCBI Accession numbers: MG518658 - MG519278).

Species richness and diversity were analyzed using a variety of alpha-diversity metrics across neotenic and metamorphic samples (Fig. 3a-d). Metamorphosis significantly reduced diversity in fecal and skin samples as follows; Chao1 and Observed OTUs were significantly lower in fecal and skin samples of the metamorphic samples as compared to neotenic samples (Unpaired Student t test, df=4, *p=0.0018, p=0005,* respectively). But Inverse Simpson and Shannon indices were not significantly different (*p=0.34, p=0.07*) for these two particular samples although Faith’s phylogenetic tree (PD) did indicate that microbiota diversity of these samples were significantly different from each other (*p=0.0022, p=0.0005,* respectively). Both Simpson and Shannon indices take into account richness and evenness in computing the metrics. Therefore, taxa with high relative abundance being heavily weighted in calculations while the indices are less sensitive to rare taxa when compared to richness only metrics ^35^. Interestingly, evenness as calculated by Simpson-E index was not significantly different between any sample pairs (*p>0.05* for all comparisons). Correspondingly, we inferred that low abundant taxa drive differences in diversity in fecal and skin samples. Both stomach and gut samples had richness and diversity that were not significantly different on all metrics between neotenic and metamorphic animals (*p>0.05;* Fig. 3a-d).

**Figure 3.**
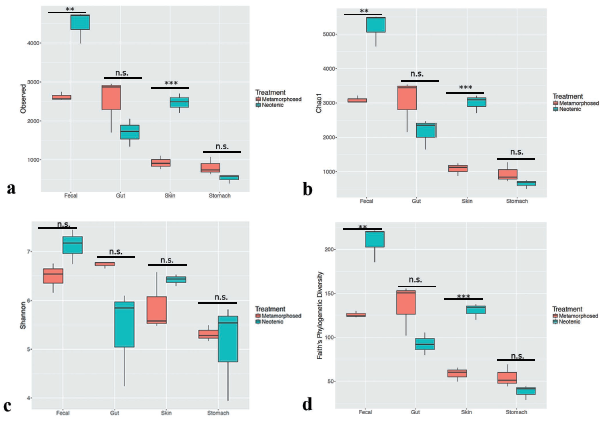
Effect of metamorphosis on bacterial diversity between neotenic and metamorphic Axolotl. Box plots illustrate the comparison of diversity indices; Observed (a), Chao1 (b), Shannon (c) and Faith’s Phylogenetic Diversity (PD) measures.

Beta diversity of bacterial communities largely differed between neotenic and metamorphic samples based on Bray-Curtis and Jaccard distance metrics (Fig. 4a; and Supplementary Fig. S2, respectively). Within and between group differences (Main-effect) using both distance metric were statistically significant as per PERMANOVA test (*Pseudo-F* (8, 18) = 7.82, *p(Monte Carlo)=0.0001*). Permutational test for homogeneity of dispersions (PERMDISP) was at the border of significance; (F (8, 18) = 8.357, *p*(perm) = 0.0507, 9999 permutations of residuals), indicating that the average within group dispersions were marginally equivalent among groups but dispersion effect, to some degree, may be confounded in the location effect. We next employed Canonical analysis of principal coordinates (CAP, Anderson and Willis, 2003) based on Bray-Curtis distance matrix, a constrained ordination that maximizes the differences among a priori groups and reveals subtle patterns, which otherwise remain elusive to unconstrained ordinations. CAP analysis clearly separated neotenic and metamorphic samples (Fig. 4b) except for the fecal samples (Correct classification rate 96.3%; trace statistics (tr(Q_m’HQ_m): 3,92174 p=0.001 with 9999 permutations, supporting rejection of the null hypothesis of no difference among the sample groups).

**Figure 4.**
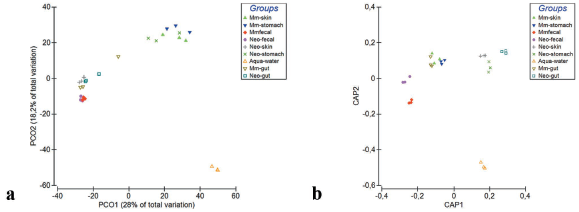
Beta diversity analysis based on Bray-Curtis distance matrix showing separation of neotenic and metamorphic bacterial communities. Samples were compared using PCO (a) and Canonical Analysis of Principal Coordinates (CAP) (b) methods.

*Firmucutes, Proteobacteria,* and *Bacteriodetes* constituted 86.7% ± 8.8 total average abundance across all Axolotl samples (Fig. 5a). Of these phyla, Proteobacteria abundance considerably increased in the skin (5.2% to 41.8%) and in the gut samples (2.7% to 9.1%). In contrast, the abundance of this phylum did not significantly change in the fecal samples yet decreased in the stomach samples (52.0% to 38.9%). Firmicutes abundance seemed to follow the opposite trend. We thus employed Pearson’s correlation to identify phylum level taxa (Supplementary Fig. S3) whose abundances negatively correlate as a result of metamorphosis. Abundances of Proteobacteria and Firmicutes along with Verrucomicrobia showed the strongest negative correlation (correlation coefficients were *r*= -0.64 and -0.54, respectively), which was also significant (False Discovery Rate (FDR) adjusted *q* =0.0027 and 0.026, respectively). Interestingly, the phylum Bacteriodetes abundance in all samples substantially increased after metamorphosis (56.7% vs. 74.2% in fecal samples; 2.5% vs. 29.7% in gut samples; 11.6% vs. 17.5% in skin samples) and negatively correlated with both Firmicutes and Proteobacteria (although this result was not significant (*q*=0.4). Overall, The Axolotl skin microbiota, among others, showed most dramatic shifts between neotenic and metamorphic stages considering these negatively correlated taxa.

**Figure 5.**
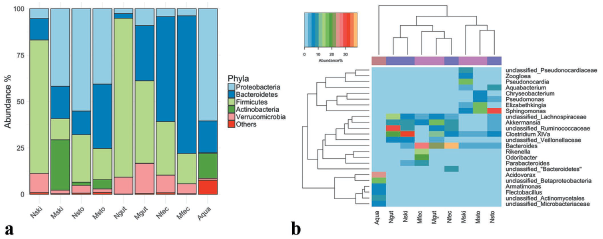
Mean relative abundances of 16S rRNA sequences. Phylum level relative abundance as bar chart (a), genus level relative abundances shown as heatmap (Individual taxa displayed if the its abundance in any sample ≥5%). Samples and bacterial taxa were clustered using average linkage hierarchical clustering of a distance matrix based on Bray–Curtis distance and taxa abundances, respectively. Samples from each group were color coded on the column side bar as follows: Aqua samples (brown); samples from neotenic Axolotl (light slate blue), metamorphic Axolotl (magenta) (b).

Percent average abundances of the genus level taxa are shown in a heatmap (Fig. 5b; for simplicity only taxa abundance ≥ 5% in any sample were shown). Samples were grouped based on Bray-Curtis similarities (top dendogram) and abundances were clustered using hierarchical clustering (average linkage). The observed differences between group similarities were driven by differences in the relative abundance of multiple bacterial taxa. For example, metamorphic skin samples clustered with stomach samples of neotenic and metamorphic Axolotl; *Clostridium IV, Bacteriodes*, and *Sphingomonas* were abundant in these samples. Notably, the genus *Elizabethkingia* were observed in high abundance in the skin and the stomach samples of the metamorphic Axolotl. Metamorphic gut sample clustered with the fecal samples; *Akkermansia and Bacteriodes* being in greater abundance in these samples. Finally, neotenic gut and skin samples were grouped together. This pattern was chiefly due to taxa that could not be classified at genus level (*unclassifiedVeillonellaceae,unclassified_Ruminococcaceae, unclassified_Lachnospiraceae*) and Clostridium XlVa. Remarkably, Aqua water samples did not have any taxa with high abundance shared with any samples from Axolotl, and separated from other samples.

### Indicator and Shared Species of Neotenic and Metamorphic Axolotl

We next used DESeq2 analysis, a negative Binomial Wald Test ^36,37^ to identify differentially abundant taxa (*q*<0.01) in neotenic and metamorphic Axolotl organs at the genus level taxonomy. We also performed indicator species analysis to identify not only abundant and rare OTUs differentially enriched in these samples but also to delineate high fidelity differential patterns (IndVal values ≥ 0.7, *p<0.01*). We largely observed concordance between both analyses. Skin samples showed the highest number of differentially enriched genera (49 taxa in metamorphic skin samples and 36 taxa enriched in neotenic skin samples); *Pseudonocardia, unclassified_Pseudonocardiaceae, Methylobacterium, Elizabethkingia, Vogesella, Chryseobacterium,* and *Zoogloea* were the top scoring genera in the metamorphic skin while *Limnohabitans* and several taxa that were classified at higher taxonomic ranking were differentially abundant in the neotenic skin samples (Fig. 6a). These genera were also detected in the indicator species analysis in several OTUs. For example, OTU89, OTU141, OTU1052, OTU1301, OTU1450, OTU2081, were classified as *Pseudonocardia* and OTU89 had the highest abundance of 15.3%. Similarly, the genus *Limnohabitans,* overrepresented in neotenic samples, was assigned to OTU1542 and OTU2587, although both OTUs had relative abundance (0.01%) that can be considered a rare OUT ^34^. The following top scoring taxa were differentially abundant in metamorphic and neotenic samples, respectively; Stomach samples: *Elizabethkingia* (OTU335) and *Limnohabitans* (OTU46); gut samples: *Elusimicrobium,* which was not detected by IndVal but OTU7777 (*Hydrogenoanaerobacterium*) was the top scoring taxa (IndVal =0.85, *P* =0.002); *unclassified_Ruminococcaceae, unclassified_Lachnospiraceae* were the indicator species of neotenic gut samples and finally *Odoribacter* (OTU14388), Rikenella (OTU14233) were both differentially abundant and indicator species of metamorphic fecal samples.

**Figure 6.**
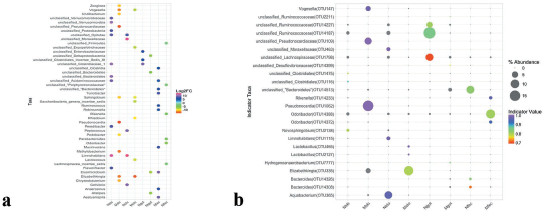
Differentially enriched genus level taxa and indicator species in samples from neotenic and metamorphic Axolotl. The color scale bar indicates log2 fold changes for absolute OTU abundances (DESeq2 analysis (*q*<0.01) (a), Bubble plot showing indicator species. Only highly significant indicator values (IndVal>0.7, *q*<0.01) are displayed. Size of bubble symbol is proportional to the mean relative abundance of indicator OTUs and the color scale bar shows indicator value for each OTU (b).

Venn diagrams showed number of OTUs shared among or unique for the samples isolated from Axolotl (Supplementary Fig. S4a and Fig. S4b). Number of unique OTUs in all samples of neotenic (7422 OTUs) and metamorphic Axolotl (6237 OTUs) was far grater than shared OTUs indicating assembly of microbiota is tissue specific as in most other animals ^38^. In terms of shared OTUs, 368 and 329 OTUs were present in all neotenic and metamorphic samples, respectively. We also collected water samples (“Aqua”) from the Aquarium of Axolotl to identify OTUs present in the water that may have colonized the Axolotl skin (Fig. 7a). We found 89 OTUs were shared by the neotenic and metamorphic skin and Aqua samples; 38 OTUs were present only in the neotenic and Aqua samples while 105 OTUs were shared by metamorphic and Aqua samples. However, relative abundances of all these OTUs were mostly less than 1%. We also compared unique and shared OTUs between gut tissue and fecal samples (Fig. 7b). Exceptionally, the number of OTUs in the metamorphic gut was 4999, representing substantial increase from 2961 OTUs found in the neotenic gut. And the two gut tissue samples shared only 227 OTUs. In stark contrast, the number of unique OTUs decreased to 1365 OTUs whereas neotenic fecal sample had 3408 unique OTUs. Notably, the shared number of fecal and gut OTUs, neotenic or metamorphic, were 288 OTUs and 272 OTUs, suggesting the fecal and gut microbiota are compositionally distinct. Finally, we identified the following genera as core taxa, i.e. shared by 90% of all samples (Core90) (Supplementary Fig. S5): *Bacteroides, Clostridium XlVa, Clostridium XlVb, Akkermansia, Odoribacter, unclassified_Veillonellaceae, unclassified_Lachnospiraceae, Parabacteroides, unclassified_Rhodospirillaceae, unclassified_Ruminococcaceae,Rikenella, unclassified_Clostridiale,* and *Desulfovibrio.*

**Figure 7.**
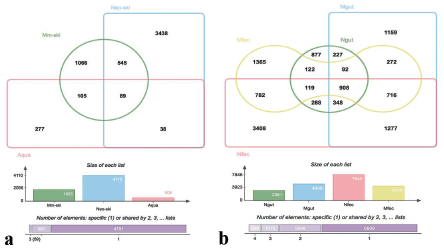
Venn diagrams showing the number of unique and shared OTUs. Skin and Aqua samples (a), and gut and fecal samples (b), collected from neotenic and metamorphic Axolotls as indicated in the diagram.

### Detection of Human taxa in Axolotl Samples and Predicted Functions

To answer the question if the captive Axolotl may have acquired human bacterial taxa we compared Axolotl microbiota with HMP stool and skin samples (HMP reference data) using weighted and unweighted Unifrac distances (Supplementary Fig. S6 (a-d)). In particular, both neotenic and metamorphic stool and skin samples showed greater similarity to HMP samples compared to other Axolotl samples. These results encouraged us to look at predicted functions of the microbiota using Phylogenetic Investigations of Communities by Reconstruction of Unobserved States (PICRUSt) Due to the specific requirement by PICRUSt, a separate OTU table was generated accordingly using a closed reference analysis based on the GreenGenes 99 database version. CAP analysis of the predicted functions revealed significant differences between the Bray-Curtis distances based on putative pathway abundances (tr(Q_m’HQ_m): 5,78539 p=0.001). However, Bray-Curtis similarities among all distinct groups were around 90%, pointing to the shared predicted functions among the bacterial microbiota of Axolotl organs (Supplementary Fig. S7).

## Discussion

The main purpose of this study was to comparatively characterize bacterial microbiota of Axolotl in neotenic and metamorphic life stages since microbiota of this important biological model has not been reported before. We also asked if differential profiles of Axolotl microbiota in the examined life stages might be accompanied by a change in host regenerative capacity. Our results show that (a)- substantial shifts occurs in the structure and composition of microbiota, particularly in the skin but also in digestive organs; and that (b)- regenerative capacity in the limbs of metamorphic Axolotl steeply diminishes while neotenic Axolotl maintains normal regenerative capacity under identical conditions and that the shifts in the skin microbiota appears to be related to the reduced limb regenerative capacity. Although the scope of this report does not include pinpointing molecular mechanisms accounting for the reduced limb regeneration and its putative association with microbiota, our observations align well with recently published experimental evidence supporting the conclusion that limb regenerative capacity of Axolotl diminishes upon metamorphosis ^33,39^. Crucially, our ongoing experiments suggest that reduced regenerative capacity is not observed for some other types of tissues in metamorphic Axolotl (Demircan *et al*., unpublished data).

We detected drastic expansion of Proteobacteria in metamorphic Axolotl skin relative to the neotenic skin and noted steep decrease of bacterial diversity in the fecal and skin samples (see Fig. 3). Evidence of skin dysbiosis and the linkage of members of Proteobacteria to reduced regenerative capacity comes from the study by ^40^; in this study pathogenic shifts in microbiota and infections were implicated in reduced regeneration in planaria, a model for tissue regeneration; Although the abundance of *Vogesella, Pseudomonas, and Sphingomonas,* all a member of Proteobacteria, and *Chryseobacterium* within Bacteroidetes phylum were virtually not detected in neotenic planarian skin samples, the abundance of these genera considerably increased in the skin of the metamorphic planarian and these genera demonstrably impeded TAK1/MKK/p38 signaling pathway of the innate immunity of the planarian host. Strikingly, our analysis revealed that abundance of these genera and some other members of Proteobacteria known to cause nosocomial infections were, too, highly significantly increased in the metamorphic skin samples from Axolotl (Fig6a and 6b); *Vogesella* (*q=*7.4E-11); *Chryseobacterium* (*q=*5.4E-12); *Undibacterium* (*q*=7.01E-11), and *Sphingomonas* (*q*=4.03E-09). The average abundance of *Pseudomonas*, too, increased (from 0.1% in neotenic skin to 2.2% in metamorphic skin) but the increase was barely significant after FDR correction for multiple testing (*q*=0.056). Furthermore, infiltration of macrophages, as key players of innate immune system, to the site of amputation and cellular signaling pathways in Axolotl were shown to play crucial role in Axolotl limb regeneration ^41^, lending further credence to the role dysbiotic microbiota can play in reducing the limb regenerative capacity by probably impeding signaling pathways of macrophages. Indeed, macrophages are crucial in removing cellular debris and regulating the balance between fibrosis and scarring ^42,43^. Metamorphosis leads to remarkable alterations in immune system such as expression of antimicrobial peptides in epidermis, increase in lymphoid Scells ratio and increasingly antigen responsive B cells, and corticosteroid-mediated apoptosis of susceptible lymphocytes ^44^. Considering the cross talk between microbes and the immune system, transformation of skin immunity (thickening mucus layer and increased secretion of antimicrobial peptides) may partly account for the disruption of microbiota after metamorphosis. Importantly, disruption in microbiota increases the risk of pathogen infection and overgrowth of opportunistic pathogens during complete metamorphosis as in the example of *Galleria mellonella*, leading to fitness costs ^8^. Taken together, the body of evidence on metamorphosing host microbe interactions support the conclusion that dysbiotic restructuring microbiota in Axolotl skin may have contributed to the reduction in limb regeneration. However, further experiments must be performed to ascertain direct mechanistic evidence supporting the cross talk between members of Proteobacteria and blastema pathways involved in regeneration. Future experiments should also include eukaryotic microbiota since fungal skin infections (e.g. chytridiomycosis) in amphibians adversely affects the vital function of amphibian skin^45,46^.

The results from this study add to the emerging appreciation for the broader roles of microbiota in regeneration and wound healing ^47-50^. Conceivably, both intrinsic (e.g. age, morphology) and extrinsic factors (e.g. diet, microbiota) may contribute to reduced limb regenerative capacity of the metamorphic Axolotl. Even though we cannot rule out other subtle intrinsic biological factors we selected adult siblings to control for the potentially confounding age factor and to minimize the genetic differences among individuals in this experiment. All animals used in our experiments were also fed on the same diet and maintained under identical conditions. Microbiota is important extrinsic factor profoundly influencing host biology particularly by interacting with the host immune system.

Our results in terms of the composition of Axolotl microbiota broadly parallel the previous reports on microbiota profile of amphibians, and salamanders in particular. Five major dominating phyla, Firmicutes, Bacteroidetes, Proteobacteria, Verrucomicrobia and Actinobacteria were abundant among all studied samples, which is consistent with previous amphibian studies ^32,51-54^. Salamander microbiota was previously studied but often distinct species of salamander skin microbiota was profiled ^51,53^ and systematic investigation of microbiota diversity in multiple organs of salamander species, particularly that of Axolotl, has been missing in the literature with few exception. Bletz *et al.* (2016) characterized both gut and skin microbiota of fire salamanders within the natural habitat. Surprisingly, like captive Axolotl in this study, the wild salamanders’ gut and skin are associated with the above mentioned five major phyla. The genera *Chryseobacterium, Pseudomonas, Flavobacterium, Sphingobacterium, Novosphingobium*, were reported to be dominant taxa in skin of fire salamander living in nature. Interestingly, we detected these genera in the neotenic skin samples in this study albeit in low abundance but substantially increased in abundance in the metamorphic skin. These bacterial genera belong to a large, ecologically diverse group, and their members include known pathogens; some could be opportunistic while some others can outcompete emerging fungal pathogens ^55^. Interestingly, our analysis revealed that metamorphic Axolotl skin was dominated by *Pseudonocardia* (15.7% ± 10.4), and an unclassified taxa from *Pseudonocardiaceae* family (9.7% ± 5.7), both are indicator species of the metamorphic skin (IndVal =0.99, *p*=0.006). *Pseudonocardia spp*. is a well known antifungal commensal microorganism ^56^ and colonize on the integument of fungus-gardening ant species. Recruitment of the members of this genus in high abundance by metamorphic Axolotl might reflect host-symobiont synergy against pathogenic fungi.

We sequenced pools of water samples from the Axolotl aquarium (“Aqua”) to identify water-borne bacterial taxa acquired by Axolotls. Surprisingly, most abundant genera of Aqua samples (e.g. *Acidovorax, Armatimonas, Flectobacillus*) were not associated with Axolotl organs but only a subset of low-abundance bacteria was detected. For example, *Aquabacterium* abundance in metamorphic skin and neotenic stomach were 3.9% and 7.3%, respectively while its abundance in Aqua sample was 0.1%. Our results are consistent with previous work reporting amphibian skin may select rare taxa from the environment ^51-53,57,58^. In this study, we found that 89 out of 509 OTUs in Aqua samples were present in neotenic skin samples while 105 OTUs shared between Aqua and metamorphic skin samples (see Fig. 6), and 150 shared OTUs with Aqua samples, the highest among others, with metamorphic stomach samples. Consequently, our results support the notion that both host and external factors shape the host microbiota but host genetics applies selective filter.

Conversely, diet is another crucial factor strongly influencing structure of gut microbiota of animal host and even dominate host genotype ^59^. Although host genotype and diet are constant in this study, metamorphosis is likely to cause remodeling of epigenetic landscape in the host genome, which in turn is expected to reshape microbiota. Notably, fecal samples from the neotenic and metamorphic Axolotls clustered together, albeit richness in metamorphic fecal samples significantly decreased. We observed that feeding behavior of the metamorphic Axolotl change during metamorphosis, the animals tend to eat less often (low appetite), which might account for the decreased fecal diversity. Although no major restructuring of intestine *via* metamorphosis was apparent as described before ^16^, we observed a higher number of goblet cells and thicker mucus layer in the metamorphic gut tissue compared to neotenic gut (Supplementary Fig. S8). Taken together, relative influence of diet and host epigenetics seems to be compartmentalized; diet appears to influence strongly the bacterial diversity in the fecal microbiota in the gut lumen whereas the host epigenetics (and the resulting changes in transcriptome due to metamorphosis) seems to play a greater and selective role in gut tissues (crypts). Some genera, often associated with symbiosis such as *Alistipes* and *Elusimicrobium,* were differentially enriched in the metamorphic gut tissues whereas neotenic gut tissues were represented by two genera with *unclassified Clostridiaceae* and *unclassified Enterobacteriaceae.* Suprisingly, the abundance of *Akkermansia* considerably increased in the metamorphic gut tissues (16%) relative to neotenic gut (9%). *A. muciniphila* within this genus is known to be a mucin degrading and symbiotic bacterium and the increase in abundance of this genus is meaningful with increasingly thicker mucus layer we observed in the metamorphic gut tissue staining.

Our analysis revealed that Axolotl microbiota to certain extent was “humanized” in captivity as manifested from similarity of gut and skin microbiota with the human microbiota, which prompted us to compare predicted functions using PICRUSt ^60^, which is optimized for human microbiota. We found that all samples included in microbiota analysis were highly similar based on the abundance of predicted microbial genes (40%). Though further study using shotgun metagenomics technology is warranted, our findings raise the possibility that deficiency in the microbial community function in a given organ can be possibly compensated without taxonomic coherence. However, caution should be exercised in interpreting these results considering the accuracy of the predicted functions is predicated on the closed reference database, whereas the majority of OTUs in this study were clustered using open reference.

## Conclusions

Our study shows microbiota inhabiting Axolotl organs considerably restructure upon metamorphosis and expansion of opportunistic bacteria within Proteobacteria may be contributing to the reduced limb regenerative capacity. These results must be taken into consideration when captive Axolotl is employed in regeneration studies. Lack of systematic studies on Axolotl microbiota has hindered its potential as a fruitful model in host-microbe interaction studies. Therefore, the data presented here make a significant contribution for further characterization of a valuable biological model for regeneration, aging, and stem cell research.

## Methods

### Ethical statement and experimental design

The local ethics committee of the Istanbul Medipol University (IMU) authorized experimental protocols and animal care conditions (the authorization number: 38828770-E.7856). All experiments were performed in accordance with relevant guidelines and regulations. The experimental design followed in this study is depicted in Fig. 1. Briefly, a total of 48 adult Axolotls (12-15 cm in length, 1 year old) were obtained from the animal care facility of the IMU. Axolotls were chosen from among the siblings. Out of 48 Axolotls 30 were reserved for amputation/limb regeneration experiments; and 18 out of 48 were used for metamorphosis experiments; half of both groups (15 for regeneration experiments and 9 for metamorphosis; 24 total) were then induced individually to undergo metamorphosis by L-thyroxine (Sigma-Aldrich, St Louis, MO, USA, Cat. No. T2376) as described elsewhere ^25^. Briefly, T4 solution was prepared by dissolving L- thyroxine in Holtfreter’s solution (final concentration 50 nM) and was administrated to animals that were maintained individually throughout the experimental period. The medium was replaced with freshly prepared T4 containing solution every third day and animals were monitored for morphological changes. Administration of the hormone continued for another 3 weeks until fully metamorphic Axolotls were obtained. Both neotenic and metamorphic animals were maintained in individual aquaria water at 18 ± 2°C in Holtfreter’s solution. Axolotls were maintained in the university animal facility within a batch-flow aquarium system where the batch water was treated with UV light and filtered to prevent infections. All animals were kept in the same aquatic solution and fed with same diet. Animals were fed once a day using a staple food (JBL Novo LotlM, Neuhofen, Germany). Axolotls did not receive any antibiotic treatment throughout the experiments.

### Sample collection

Animals were sacrificed using 0.2% MS222 (Sigma-Aldrich, St Louis, MO, USA. Cat. No. E10521) approximately in two months upon visual observation of metamorphosis. Neotenic and metamorphic animals formed two major experimental groups. Randomly selected three animals were grouped to have three replicates (R1, R2 and R3) for neotenic and metamorphic experimental groups (Fig. 1). For each replicate skin, stomach, intestine and fecal samples were harvested from three animals and pooled together under sterile conditions in a bio safety cabin. To collect the skin samples, animals were rinsed in sterile water to get rid of the transient bacteria. As the next step, skin samples from the mid stylopod level of the right forelimb were isolated for each replicates with punch biopsy using disposable and sterile 6 mm diameter punches (Miltex, York, PA, USA). Stomach and intestine samples were collected after dissection of animals. Intestine contents were first removed, split into approximately equal 4-5 pieces, and rinsed with sterile serum physiologic solution five times. Fecal samples were harvested from the rectum. Isolated and pooled samples for each replicate were frozen in liquid nitrogen immediately for cryopreservation. All samples were stored at -80°C till DNA isolation. To compare harvested samples with the microbial structure of Holtfreter’s solution (“Aqua” samples), water samples from randomly chosen aquariums (n=9) were collected and 3 samples were pooled together to have three replicates (R1, R2 and R3).

### Amputation

A total of 30 Axolot were used for regeneration experiments. Half of the animals were induced to undergo metamorphosis by T4 administration as described above. After complete metamorphosis was achieved, right forelimb of both neotenic and metamorphic animals were amputated from the mid-stylopod site Both macroscopic and microscopic pictures were taken by using Nikon D3200 camera and Zeiss Axio zoom V16 microscope, respectively. Animals were anesthetized in 0.1% MS222 (Sigma-Aldrich, St Louis, MO, USA. Cat. No. E10521) for all animal procedures.

### Histology

Removal of fecal from the isolated intestine samples was followed by fixation in 10% neutral buffered formalin (NBF) for 48 h. The samples were then processed by immersion of materials in ascending alcohol series, toluene and embedding in paraffin. 4 µm thick tissue sections were obtained by using microtome. Sections were deparaffinized and stained with Hematoxylin and Eosin (Bio-Optica Mayer’s Hematoxylin and Eosin Y Plus), Masson’s Trichrome (KIT, Masson Trichrome with aniline blue, Bio Optica, 04- 010802), Alcian Blue (KIT, Alcian Blue Acid Mucopolusaccharides staining, Bio-Optica, 04-160802) according to manufacturer’s instructions. Images were taken by using the NIKON DS-Fi2-U3 Digital Camera. The detailed protocol was described in a previous work ^16^.

### DNA extraction

DNA isolation from the skin, intestine, and stomach samples was carried out with DNeasy Blood & Tissue kit (Qiagen) according to the manufacturer’s protocol. QIAamp DNA Stool Mini Kit (Qiagen) was used to extract the DNA of the fecal samples by following the manufacturer’s recommendations. To extract DNA from the water samples were first filtered through filters with 0.2 μm pore size and the filter papers subsequently were used in DNA extractions using metagenomic DNA Isolation Kit for Water (Epicentre, Cat. No. MGD08420) by following the producer’s protocol. Spectramax i3 (Molecular Devices, Sunnyvale, CA) was used to measure the concentrations of isolated DNA. Quality of DNA samples was checked by electrophoresis in 1.0% agarose gels.

### PCR and sequencing of 16S rRNA amplicons

To amplify the variable V3-V4 regions of the 16S rRNA gene, the primers 341F (5′- CCTACGGGNGGCWGCAG -3′) and 805R (5′- GACTACHVGGGTATCTAATCC-3′) were used ^61^. MiSeq sequencing adaptor sequences were added to the 5′ ends of forward and reverse primers. Approximately 12.5 ng of purified DNA from each sample was used as a template for PCR amplification in 25 μl reaction mixture by using 2x KAPA HiFi HotStart ReadyMix (Kapa Biosystems, MA, USA). For PCR amplification, the following conditions were followed: denaturation at 95° C for 3 min., followed by 25 cycles of denaturation at 95°C for 30 sec., annealing at 55°C for 30 sec. and extension at 72°C for 30 sec., with a final extension at 72°C for 5 min. No template negative control samples were included to check PCR contamination and none of the negative controls yielded detectable level of amplification on agarose gels. Amplified PCR products were purified with Agencourt AMPure XP purification system (Beckman Coulter) and Nextera PCR was performed by using sample-specific barcodes. Constructed Nextera library was then sequenced by Illumina MiSeq platform using MiSeq Reagent Kit v3.

### Sequence processing, clustering, and taxonomic assignment

To analyze the paired-end sequencing data Quantitative Insights Into Microbial Ecology (QIIME, v1.9.1) ^62^ software was used at the Nephele platform (v.1.6, 2016) of the National Institute of Allergy and Infectious Diseases (NIAID, Bethesda, MD). Nephele platform was also used for the Phylogenetic investigation of communities by reconstruction of unobserved states (PICRUSt) analysis ^60^ and for comparing the data with the Human Microbiome Project (HMP). Before submitting the raw reads into the Nephele pipeline, primers were removed using cutadapt program ^63^ and the pipeline was stringently configured to perform the following steps; reads below average quality scores (q<30) and read-length >450 bp were eliminated. After joining the pair-end reads, the reads were clustered into operational taxonomic units (OTUs) using open reference OTU- picking strategy ^64^. The open-reference approach initially runs a closed-reference step. Sequences that fail closed-reference assignment are then clustered as *de novo* OTU based on pairwise similarity among all sequences in the data set. Open reference clustering was performed based on the 97% clustered SILVA reference (SILVA 123 release;) database ^65^ and SortMeRNA combined with SUMACLUST algorithms ^66^. Non-matching reads to closed reference were subsequently clustered *de novo.* After obtaining the OTU table from the pipeline, UCHIME (v.4.2) program (http://drive5.com/uchime) integrated into the Mothur (v.1.39.5) tool ^67^ was separately run to remove chimeric reads. Additionally, The RDP classifier ^68^ (v. 2.2) was locally run to assign taxonomy for each OTU at a confidence greater than 70% cutoff. Reads that could not be classified at the genus level were sequentially assigned to higher taxonomic hierarchy up to the kingdom level. Unclassified reads at the kingdom level (“Unclassified Bacteria”) were extracted from the OTU-representative sequences and searched for nearest neighbor method using MOLE- BLAST ^69^. This tool computes a multiple sequence alignment (MSA) between the query sequences along with their top BLAST database hits, and generates a phylogenetic tree. Species richness and diversity were estimated by QIIME with the following alpha diversity metrics: OTU richness, Chao1, Shannon, Simpson E, Inverse Simpson, and Faith’s phylogenetic diversity (PD). We assessed normality of alpha diversity data using Shapiro-Wilk tests and compared the metrics between neotenic and metamorphic microbiota using unpaired two-tailed *t*-test. Venn diagrams were constructed using jvenn, a web based tool (http://jvenn.toulouse.inra.fr) ^70^.

### Multivariate analysis of community structures and diversity

Bray–Curtis similarity index ^71^ and Jaccard index of similarity ^72^ were used to obtain distance matrix after standardizing by the column sums and transforming (square-root) the read abundance data. Similarities in microbial community structures among samples were first displayed using principal coordinate analysis (PCO) (unconstrained). Differences in community structure related to metamorphosis were displayed using a constrained ordination technique, Canonical Analysis of Principal coordinates (CAP). Tests of the multivariate null hypotheses of no differences among a priori defined groups were examined using PERMANOVA and the CAP classification success rate. CAP uses PCO followed by canonical discriminant analysis to provide a constrained ordination that maximizes the differences among a priori groups and reveals patterns that cannot be unraveled using unconstrained ordinations ^73^. CAP classification success rates and CAP traceQ_m0HQ_m statistics were examined in combination to draw conclusions about separation of a priori groups. Permutational analysis of multivariate dispersions (PERMDISP) ^74^ was used to test for heterogeneity of community structure in a priori groups. PERMANOVA, CAP and PERMDISP were performed with 9999 permutations and run as routines in PRIMER6 ^75^.

To delineate bacterial taxa responsible for the multivariate patterns and differentially enriched taxa between the neotenic and metamorphic Axolotl organs, we used DESeq2, a negative Binomial Wald Test ^36,37^ and indicator species analysis ^76^. DESeq2 analysis results, along with core OTU heatmap, phylum correlation heatmap, and read count figures were obtained using MicrobiomeAnalyst ^77^. To perform indicator species analysis, R package labdsv was used. Boxplots, barchart, bubbleplots, and heatmaps were generated using R packages including vegan, ggplot2, heatmap2, Heatplus, reshape, colorramps, and RcolorBrewer (R Core Team 2016, http://www.r-project.org).

### Availability of data and materials

The datasets generated and/or analysed during the current study are available in the figshare repository: https://figshare.com/s/b1f232d8d054e0e20f06

## Acknowledgements

This study was financially supported by the Medipol University Research Fund. This study used the Nephele platform from the National Institute of Allergy and Infectious Diseases (NIAID) Office of Cyber Infrastructure and Computational Biology (OCICB) in Bethesda, MD.

## Author contributions statement

TD and SY conceived the study. TD, BY, İK, AEİ and ECF performed animal experiments. TD, GÖ, and SY acquired sequencing data; TD, GO, and SY analyzed and interpreted the data. TD, GO, GÖ, and SY drafted and critically reviewed/revised the manuscript. All authors read and approved the final manuscript.

